# Multiple steps of prion strain adaptation to a new host

**DOI:** 10.1101/2023.10.24.563743

**Authors:** Olga Bocharova, Natallia Makarava, Narayan P. Pandit, Kara Molesworth, Ilia V. Baskakov

**Author notes:** These authors contributed equally to this work (Olga Bocharova, Natallia Makarava). Address correspondence to: Ilia V. Baskakov, Center for Biomedical Engineering and Technology, University of Maryland School of Medicine, Baltimore, 111 S. Penn St., Baltimore, MD, 21201, USA.

## Abstract

The transmission of prions across species is a critical aspect of their dissemination among mammalian hosts, including humans. This process often necessitates strain adaptation. In this study, we sought to investigate the mechanisms underlying prion adaptation while mitigating biases associated with the history of cross-species transmission of natural prion strains. To achieve this, we utilized the synthetic hamster prion strain S05. Propagation of S05 using mouse PrP^C^ in Protein Misfolding Cyclic Amplification did not immediately overcome the species barrier. This finding underscores the involvement of factors beyond disparities in primary protein structures. Subsequently, we performed five serial passages to stabilize the incubation time to disease in mice. The levels of PrP^Sc^ increased with each passage, reaching a maximum at the third passage, and declining thereafter. This suggests that only the initial stage of adaptation is primarily driven by an acceleration in PrP^Sc^ replication. During the protracted adaptation to a new host, we observed significant alterations in the glycoform ratio and sialylation status of PrP^Sc^ N-glycans. These changes support the notion that qualitative modifications in PrP^Sc^ contribute to a more rapid disease progression. Furthermore, consistent with the decline in sialylation, a cue for “eat me” signaling, the newly adapted strain exhibited preferential colocalization with microglia. In contrast to PrP^Sc^ dynamics, the intensity of microglia activation continued to increase after the third passage in the new host. In summary, our study elucidates that the adaptation of a prion strain to a new host is a multi-step process driven by several factors.

## Introduction

Prion diseases encompass a group of invariably fatal transmissible neurodegenerative conditions affecting both humans and other mammals (1). The transmissible agent responsible for prion diseases consists of a prion protein in a self-propagating state rich in β-sheets, known as PrP^Sc^ (2–4). Prions replicate by enlisting and converting the normal, cellular form of the same protein (PrP^C^) into disease-associated states. Within the same host, prions induce various disease phenotypes, often characterized by differing cell tropism, affected brain regions, patterns of PrP^Sc^ deposition, and incubation periods leading to disease (5–8). The multitude of disease phenotypes is attributed to a structural diversity of PrP^Sc^ states, referred to as prion strains that are formed within the same amino acid sequence of PrP (5,9,10). According to the cloud hypothesis, individual prion strains are structurally heterogeneous too and consist of major and minor variants (11).

PrP^C^ undergoes posttranslational modifications, involving one or two sialylated N-linked glycans, along with a glycosylphosphatidylinositol (GPI) anchor (12–15). Although cells express a multitude of PrP^C^ sialoglycoforms, different prion strains exhibit varying degrees of selectivity in recruiting these PrP^C^ sialoglycoforms (16–18). For instance, hamster strains such as 263K or Hyper do not exhibit specific preferences for particular glycoforms, as the glycoform ratio in their PrP^Sc^ mirrors that of PrP^C^ (17,19,20). Conversely, mouse-adapted strains including RML, 22L, and ME7, display strong preferences and selectively recruit hyposialylated and monoglycosyled PrP^C^ molecules (17,19,20). While individual strains specify a subset of sialoglycoforms that can fit into their structure, the presence or absence of N-glycans at one or both sites can provide a selective advantage for the replication of certain prion strains. PrP^Sc^ strains could be selectively amplified from a mixture in Protein Misfolding Cyclic Amplification by manipulating the glycosylation status of PrP^C^ supplied to the reaction (21).

The success of prion transmission between different species is often significantly lower than the transmission within the same host, primarily due to the presence of a species barrier (22–24). The species barrier manifests itself as prolonged incubation time to disease, a low attack rate, or a lack of clinical disease in the initial passages (25,26). Following cross-species transmission, prion strains often adapt to a new host upon serial passaging (25,27). Among common manifestations of strain adaptation are an increase in attack rate, a reduction of the incubation time to disease, and changes in the disease phenotype (27,28). To a large extent, the species barrier is believed to be attributed to differences in amino acid sequence between host PrP^C^ and donor PrP^Sc^, also known as a sequence barrier (29–34). Predicting the strength of the species barrier based solely on differences in amino acid sequences is a challenging task, emphasizing the significant role played by other contributing factors. Among these factors are the strain-specific characteristics of the host’s PrP^Sc^ and the route of transmission, which also have a substantial influence on the species barrier and the outcome of strain adaptation (33,35–39). Additionally, post-translational modifications of PrP^C^ including its glycosylation and sialylation status also contribute to shaping the species barrier (16,17,40).

To shed light on prion adaptation, the majority of prior investigations relied on prion strains of natural origin. These prions were isolated from domesticated or wild animals and subjected to serial passaging through various rodent species (41–46). For example, the widely employed hamster-adapted strain 263K (also known as Sc237), which initially originated from goats, underwent successive passaging in mice and rats, before finally adapting to Syrian hamsters (44). According to the cloud and deformed templating hypotheses (11,47,48), cross-species transmission alters the composition of PrP^Sc^ variants within a strain and may even give rise to novel variants. Predicting the extent to which a strain’s history of passaging through different mammalian species affects its behavior in crossing the species barrier along with the outcomes of adaptation to a new species is challenging.

To investigate the mechanisms of prion adaptation without the biases introduced by a history of cross-species transmission inherited to strains of natural origin, we employed a prion strain of synthetic origin known as S05. S05 was initially generated in Syrian hamsters through the inoculation of amyloid fibrils produced from hamster recombinant PrP *in vitro* (49). Unlike hamster-adapted strains of natural origin, S05 has never undergone passaging in species other than Syrian hamsters. To eliminate the sequence barrier, the hamster strain S05 was first adapted to the mouse PrP^C^ sequence using serial Protein Misfolding Cyclic Amplification with beads (sPMCAb) before transmission to the new host. Despite successfully overcoming the sequence barrier, the subsequent adaptation to mice was found to be a time-consuming process that required at least five serial passages. This extended process of strain adaptation allowed us to dissect mechanistic details and shed new light on the phenomenon of strain adaptation.

## Results

The direct transmission of the hamster S05 strain to C57Bl/6J mice via the intracerebral (IC) route proved unsuccessful in inducing clinical disease. Moreover, no traces of PrP^Sc^ were detectable in the mice challenged with S05 even at 482 days post-inoculation (dpi) (Figure 1B). These findings clearly highlight a significant species barrier preventing the successful transmission of the S05 strain to mice. It is anticipated that replication of a prion strain *in vitro* using PrP^C^ substrate from the new host could potentially diminish or entirely abrogate this species barrier (50,51). In order to adapt the hamster S05 strain to a mouse (Mo) PrP sequence, brain-derived S05 was employed to seed serial protein misfolding cyclic amplification with beads (sPMCAb) conducted in mouse normal brain homogenate (BH) (52). Hamster S05 showed steady amplification in Mo sPMCAb (Figure 1A). This observation suggests that the specific PrP^Sc^ structures unique to the S05 strain effectively facilitated the recruitment and conversion of Mo PrP^C^ in an *in vitro* environment.

**Figure 1.**
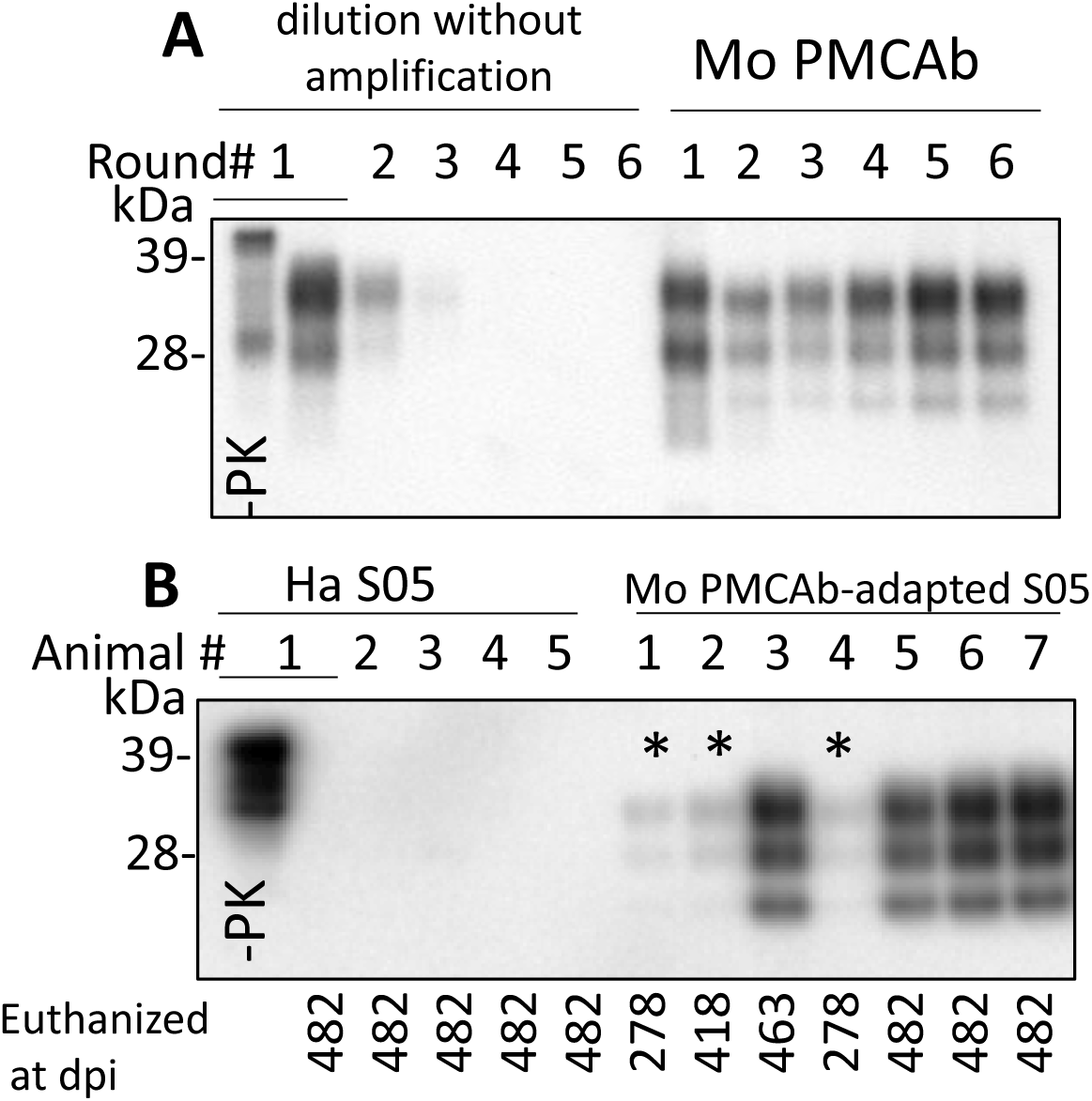
Adaptation of hamster S05 strain to mouse PrP^C^. (**A**) For adapting the hamster S05 strain to mouse PrP^C^, sPMCAb was seeded with 10^3^-fold diluted S05 BHs and carried out using mouse normal BH as a substrate. Corresponding serial dilutions of S05 BHs in the absence of amplification are shown as references. Immunodetection with SAF-84. (**B**) Western blot analysis of PrP^Sc^ in mice inoculated IC with 10% of S05 BH or mouse sPMCAb-adapted S05. Products of the tenth sPMCAb round were used for inoculation. Animals # 1, 2 and 4, marked by asterisks, were euthanized for testing the presence of PrP^Sc^ at the early stages. Immunodetection with ab3531. In **A** and **B,** all samples, with the exception of lanes indicated as -PK were treated with PK.

C57Bl/6J mice challenged via IC route with Mo sPMCAb-adapted S05 did not exhibit any clinical signs of disease in the first passage. However, mice euthanized at 463 and 482 days post-inoculation (dpi) displayed notable PrP^Sc^ signals on Western blots (Figure 1B). Even mice euthanized at earlier time points (278 dpi and 418 dpi) showed faint PrP^Sc^ bands (Figure 1B). In summary, all seven mice inoculated with Mo sPMCAb-adapted S05 exhibited PrP^Sc^ in their brains. In contrast, all five mice challenged with hamster brain-derived S05 tested negative for PrP^Sc^. Despite the absence of clinical disease in mice inoculated with Mo sPMCAb-adapted S05, these findings demonstrate that the adaptation of S05 to Mo PrP in sPMCAb contributed to a reduction in the species barrier, although complete abolition was not achieved.

To adapt S05 to mice, a serial transmission of brain-derived PrP^Sc^ from the mouse challenged with Mo sPMCAb-adapted S05 and euthanized at 482 dpi was conducted (Figure 2). In the second passage, all the mice exhibited noticeable clinical signs, such as clasp-related issues, impaired mobility, and a rigid tail. Additionally, there was a noticeable accumulation of PrP^Sc^ in their brains (Figure 2C). The incubation times to terminal disease exhibited a broad range spanning from 278 to 370 dpi (Figure 2A, B). This variation suggests that the strain had not yet fully adapted to its new host. In the third passage, the incubation time to terminal disease dropped significantly to 152±1.8 dpi relative to 311±20.4 dpi seen in the second passage (Figure 2A, B), and the animal cohort progressed to the terminal disease synchronously. Surprisingly, the trend of decreasing incubation time persisted in the fourth and fifth passages, reaching 132±3.3 dpi and 124±0.8 dpi, respectively (Figure 2A, B). Although the reduction in the incubation time during the fourth and fifth passages was moderate, the differences between the third and fourth, as well as the fourth and fifth passages, proved to be statistically significant (Figure 2B). The serial transmission experiment yielded noteworthy insights, indicating that even after adapting to the Mo PrP amino acid sequence, at least five sequential passages were necessary to achieve adaptation of the S05 strain to mice.

**Figure 2.**
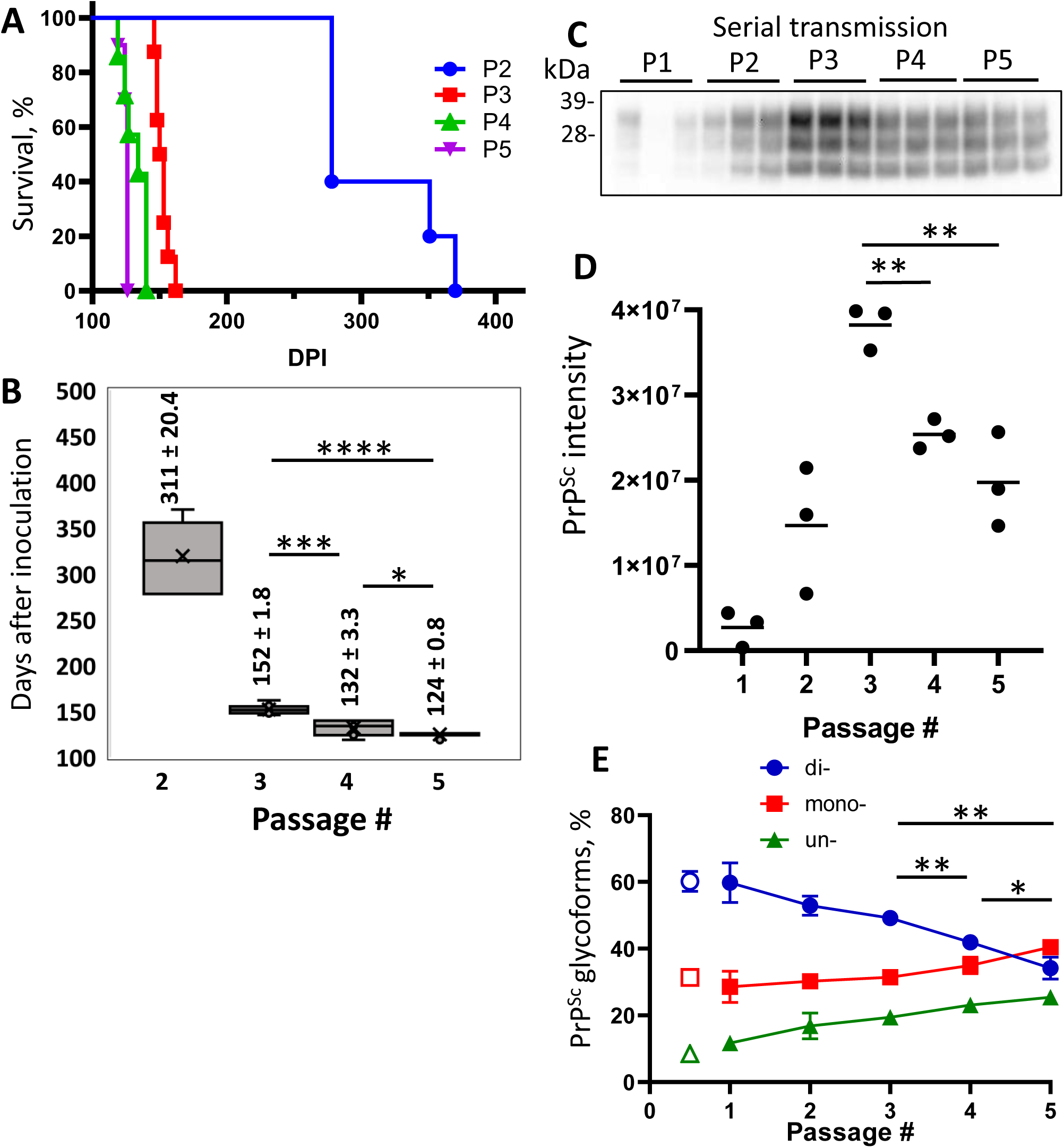
Adaptation of S05 to mice. Mice were inoculated IC with Mo sPMCAb-adapted S05 (P1) and then serially passaged four times (P2-P5). (**A**) Kaplan-Meier survival plot for the serial passages (P2 to P5) of Mo sPMCAb-adapted S05 in mice. (**B**) Box-and-whisker plot of incubation time to terminal disease in mice as a function of serial passage number. The midline of the box- and-whisker plot denotes the median, the x represents the mean, and the ends of the box plot denote the 25^th^ and 75^th^ percentiles (n=5 in 2^nd^, n=8 in 3^d^, n=7 in 4^th^ and n=10 in 5^th^ passages). Data presented as means ± SD, *P≤0.05, ***P≤0.001, ****P≤0.0001, two-tailed unpaired t-test. (**C**) Western blot analysis of PrP^Sc^ in five passages of Mo sPMCAb-adapted S05. All samples are 10% BHs, treated with PK and detected using D18 antibody. (**D**) Quantification of PrP^Sc^ amounts in five serial passages of Mo sPMCAb-adapted S05 in mice using densitometry of Western blots. (**E**) Change in the percentage of di-, mono-, and unglycosylated PrP^Sc^ as a function of serial passage (filled symbols), and the percentage of PrP^Sc^ glycoforms in the original hamster S05 strain (empty symbols). Mean ± SD (n=3). In **D** and **E**, *P≤0.05, **P≤0.01, ***P≤0.001 by two-tailed unpaired t-test.

To investigate the dynamics of PrP^Sc^ during the course of S05 adaptation, brain materials from five consecutive passages were subjected to analysis via Western blotting. Up until the third passage, increased amounts of PrP^Sc^ were produced by the terminal stage of the disease with each subsequent passage (Figure 2C, D). Considerably higher amounts of PrP^Sc^ in the third relative to the second passage despite a halved incubation period to the disease argues that the rate of PrP^Sc^ deposition accelerates as the strain undergoes adaption to its new host. Intriguingly, the fourth and fifth passages revealed diminishing amounts of PrP^Sc^ relative to the third passage (Figure 2C, D). This outcome implies that beyond the third passage, the reduction in incubation time was no longer driven by an acceleration in PrP^Sc^ deposition rate. Rather, it likely stems from qualitative changes within the PrP^Sc^.

To investigate qualitative changes within PrP^Sc^, we examined the dynamics of glycoform ratios. In the first passage, the predominant PrP^Sc^ isoform was diglycosylated, and the proportion of PrP^Sc^ glycoforms closely resembled that of the original hamster S05 strain (Figure 2E). However, over subsequent passages, the proportions of monoglycosylated and unglycosylated isoforms gradually increased at the expense of the diglycosylated isoforms (Figure 2E). Remarkably, this trend persisted even beyond the third passage. By the fifth passage, monoglycosylated isoforms surpassed the diglycosylated isoforms in prevalence (Figure 2E). When compared to other mouse-adapted strains (SSLOW and 22L), it became evident that by the fifth passage, the glycoform ratio of Mo S05 had become characteristic of that observed in other mouse-adapted strains (Figure 3A, B). The strain-specific ratios of PrP^Sc^ glycoforms arise due to differences in the selective recruitment of diverse PrP^C^ sialoglycoforms, a process that unfolds in a strain-specific manner (17). Alterations in the glycoform ratio serve as a testament to the evolving selectivity of recruitment across five consecutive passages.

**Figure 3.**
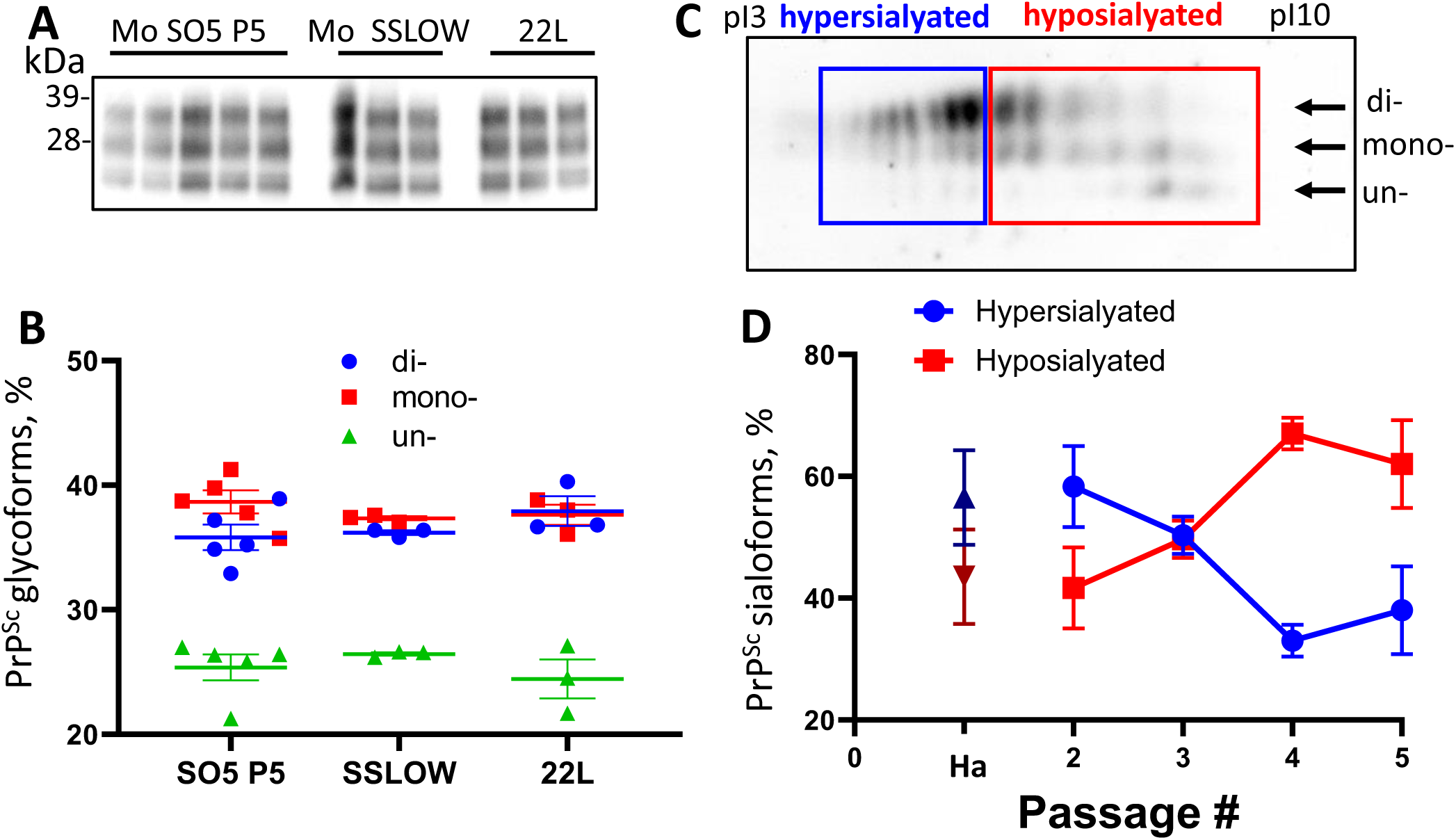
Analysis of PrP^Sc^ glycosylation and sialylation ratios. (**A**) Western blot analysis of PrP^Sc^ in Mo S05 (5^th^ passage), Mo SSLOW and 22L animals. All samples are 10% BHs, treated with PK and detected using D18 antibody (**B**) The percentage of di-, mono-, and unglycosylated PrP^Sc^. Mean±SEM (n=3 for Mo SSLOW and 22L, and n=5 for Mo S05). (**C**) An example of a 2D Western blot of Mo S05 PrP^Sc^ showing the division of sialoglycoforms into hypo- and hypersialylated isoforms. Western blot was stained with D18 antibody. Arrows indicate the position of di-, mono- and unglycosylated sialoglycoforms. (**D**) Change in relative populations of hypo- and hypersialylated isoforms in 2^nd^ to 5^th^ passages of Mo S05 and relative population of isoforms in the original S05 strain in hamsters (Ha). Mean ± SD (n=3).

A shift in selectivity of recruitment of PrP^C^ sialoglycoforms seen in the course of S05 adaptation is expected to alter a pattern of carbohydrate epitopes of PrP^Sc^ N-glycans. To test whether this is the case, a composition of PrP^Sc^ sialoglycoforms was analyzed using 2-dimensional electrophoresis followed by Western blot (3C). Because the samples are denatured prior to the 2D analysis, the distribution of charge isoforms in the horizontal dimension reflects the sialylation status of individual PrP molecules, where hyposialylated molecules run toward basic pH and hypersialylated toward acidic pH (53). On 2D, the sialoglycoforms were divided into two groups: hypersialylated and hyposialylated (Figure 3C). In the first passage, the amount of PrP^Sc^ material was not sufficient to be detectable on the 2D. In the second passage, the composition of PrP^Sc^ sialoglycoforms was found to be very similar to that of the original hamster S05 strain, where hypersialylated glycoforms dominated over hyposialylated glycoforms (Figure 3D). Akin to most of the hamster strains, the original S05 strain recruits PrP^C^ sialoglycoforms proportionally to their expression levels, which results in a vast majority of PrP molecules being diglycosylated and heavily sialylated. However, by the fourth passage of S05 in mice, the ranking order of PrP^Sc^ sialoglycoforms reversed, and the percentage of hyposialylated glycoforms dominated over that of hyposialylated glycoforms (Figure 3D). This result illustrates that (i) selectivity of recruitment changes with strain adaptation, and (ii) mouse-adapted S05 acquired a strong preference for recruiting hyposialylated PrP^C^ sialoglycoforms. To summarize, the analysis of glycosylation and sialylation status confirmed qualitative changes within PrP^Sc^.

In the course of S05 adaptation, the sialylation status of PrP^Sc^ declines resulting in the exposure of galactose residues instead of sialic acid at the terminal positions of N-linked glycans. This exposed galactose serves as the ‘eat me’ signal for microglia (54,55). Therefore next, we performed co-immunostaining for PrP^Sc^ with the markers of microglia (Iba1), astrocytes (GFAP), and neurons (MAP2) to determine whether PrP^Sc^ co-localize with microglia, astrocytes, or neurons, respectively. This analysis was performed in the cortex and thalamus regions (Figure 4A). In the cortex, a significant majority of PrP^Sc^ puncta showed co-localization with microglia, while a notably smaller proportion of PrP^Sc^ was observed in association with astrocytes (Figure 4A, B). Very limited, if any, co-localization was detected with neurons (Figure 4A, B). A similar pattern was observed in the thalamus (Figure 4A, B). Given that microglia do not express PrP^C^ and do not replicate PrP^Sc^, PrP^Sc^ puncta associated with microglia are attributed to phagocytosed materials. In summary, our co-immunostaining results demonstrate that the majority of PrP^Sc^ aggregates co-localize with microglia.

**Figure 4.**
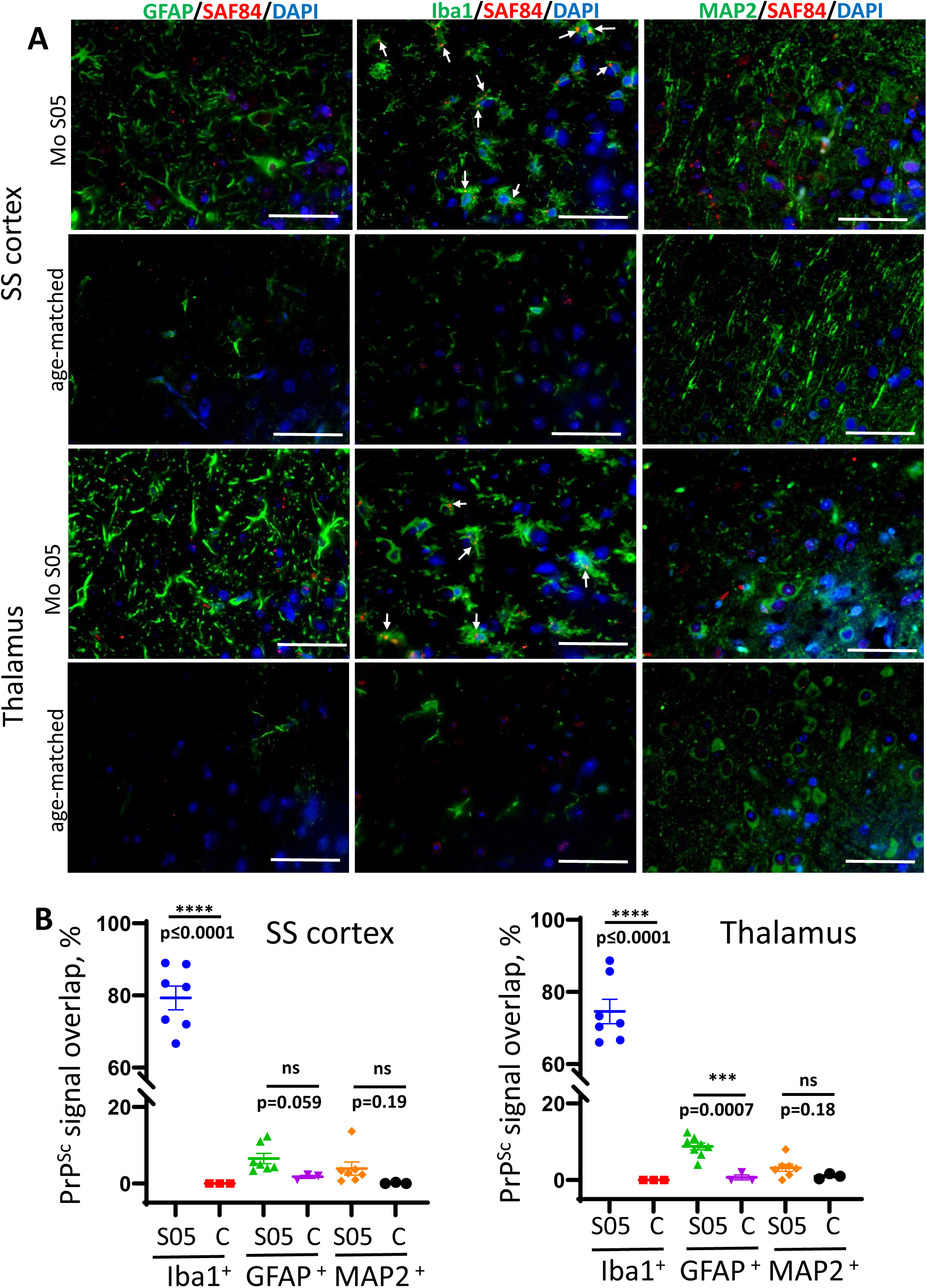
Co-localization of PrP^Sc^ and microglia, astrocytes, and neurons. (**A**) Representative images of co-immunostaining for PrP^Sc^ (SAF-84 antibody, red) with astrocytes (anti-GFAP, green), microglia (anti-Iba1, green), or neurons (anti-MAP2, green) in somatosensory (SS) cortex and thalamus in mice from the 5^th^ passage of Mo S05 or age-matched control mice. Arrows point to small PrP^Sc^ deposits colocalized with microglia. Scale bars = 50 μm. (**B**) The percentage of PrP^Sc^ signal overlapping with Iba1^+^ microglia, GFAP^+^ astrocytes, and MAP2^+^ neurons in animals from the 5^th^ passage of S05 and age-matched control mice (designated as C). Mean ± SEM (n= 7 fields of view from 3 animals per experimental group; n= 3 fields of view from 3 animals per non-infected control group).

Previous results have revealed a faster progression of the disease in the fourth and fifth passages, despite lower amounts of PrP^Sc^. This suggests that qualitative changes within PrP^Sc^ drive the second stage of adaptation. Among such changes are a gradual transformation in the glycoform ratio and the emergence of hyposialylated PrP^Sc^ with serial passaging of S05 in mice. Hyposialylated PrP^Sc^ was found in colocalization with microglia, supporting the hypothesis that reduced sialylation of N-glycans triggers the phagocytic uptake of PrP^Sc^. Previous studies have shown that the sialylation status of PrP^Sc^ defines the response of microglia (56), so next, we analyzed the dynamics of microglia and astrocyte activation during the course of S05 adaptation to mice. As judged from staining for Iba1 (Figure 5A, B), the degree of microglia activation exhibited gradual, incremental changes with serial passaging, showing a steeper uptrend after the third passage. Staining for GFAP indicated that astrocyte activation reached similar levels by the terminal stage, regardless of serial passages, except for the first passage when animals were euthanized in the absence of clinical signs. In summary, these results illustrate that qualitative changes in PrP^Sc^ are accompanied by the degree of microglial activation and suggest that neuroinflammation might serve as a more accurate indicator of terminal disease progression than PrP^Sc^ quantities.

**Figure 5.**
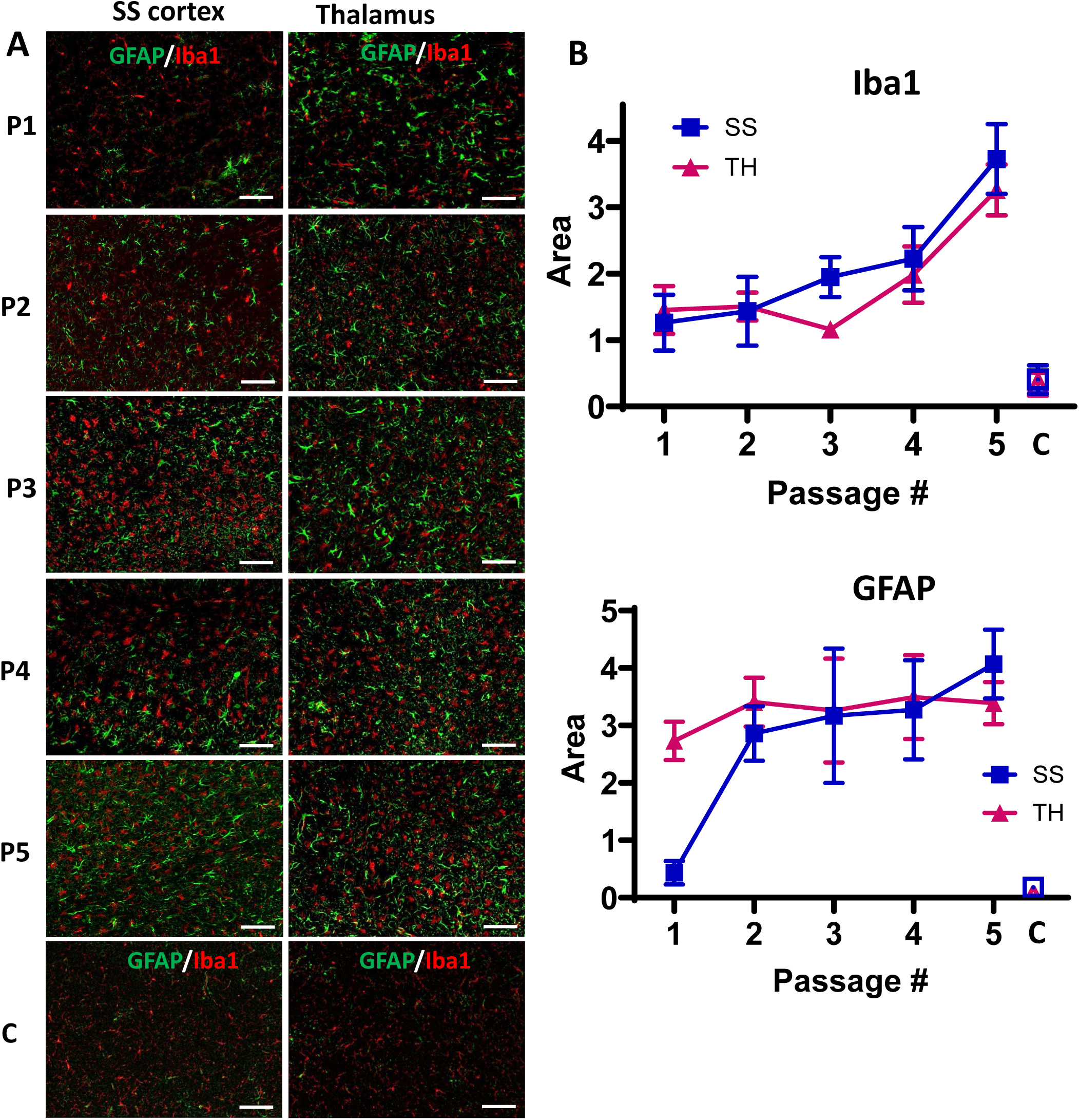
Analysis of neuroinflammation during the course of S05 adaptation to mice. (**A**) Representative images of somatosensory (SS) cortex and thalamus (TH) in mice from five serial passages of S05, and non-infected control mice (designated as C) co-immunostained with anti-GFAP (green) and anti-Iba1 (red) antibodies. The age of control mice matched the age of mice from the 4^th^ and 5^th^ passages. Scale bar = 50 µm. (**B**) Quantification of areas covered by GFAP^+^ or Iba1^+^ cells in somatosensory cortex (SS, squares) and thalamus (TH, triangles) in mice from five serial passages of S05, and non-infected control mice (C, open squares and triangles). Mean ± SD (n=7-9 fields of view from 3 animals per group).

The current study illustrates that the adaptation of a prion strain to a new host is a multi-step process driven by several factors. During the first stage of adaptation in mice, the replication of donor PrP^Sc^ using the PrP^C^ of a new host triggers alterations in PrP^Sc^ topology. When the PrP^Sc^ topologies of the donor and recipient hosts are substantially different, the new topology influences the selectivity of the recruitment of PrP^C^ sialoglycoforms. The selection of a different set of sialoglycoforms leads to changes in the decoration of PrP^Sc^ by carbohydrate moieties, ultimately resulting in alterations in the disease phenotype.

## Discussion

The present study has revealed that the adaptation of a prion strain to a new host is a multi-step process, which is driven by several factors. Notably, propagation of the hamster strain S05 using a mouse PrP^C^ substrate in sPMCAb fell short of surmounting the species barrier. This outcome underscores the involvement of factors beyond mere disparities in the primary structures of hamster and mouse PrPs. Upon consecutive transmissions in mice, a marked reduction in the incubation time to disease was observed in the second and third passages, concomitant with escalating levels of PrP^Sc^ in both passages. The increased accumulation of PrP^Sc^, alongside with abbreviated incubation times, suggests that the adaptation of the strain to a new host is driven by an acceleration in the rate of PrP^Sc^ replication. Interestingly, in the fourth and fifth passages, the disease progressed even more rapidly compared to the third passage. Unexpectedly, these shortened incubation times coincided with diminished quantities of PrP^Sc^ in each successive passage. This observation highlights that, in the fourth and fifth passages, the terminal disease state is reached with lower amounts of PrP^Sc^. This phenomenon strongly implies that, with strain adaptation, PrP^Sc^ undergoes qualitative alterations that evoke a more potent biological response, be it in terms of direct neuronal toxicity or neuroinflammation. These qualitative changes in PrP^Sc^ encompass a transformation in glycoform ratio and sialylation status. During strain adaptation, PrP^Sc^ shifts from being predominantly diglycosylated to primarily monoglycosylated. This transformation is accompanied by a reduction in sialylation levels. Notably, reduced sialylation exposes galactose residues at the terminal position of N-linked glycans, which are recognized as “eat me” signals for microglia (54,55,57,58). Consequently, the transformation of glycosylation and sialylation status bears substantial implications for the biological activity of PrP^Sc^.

Previous studies have demonstrated that the removal of sialic acid residues from PrP^Sc^ N-linked glycans enhances the pro-inflammatory response of primary microglia cultured *in vitro* (56). Moreover, PrP^Sc^ infectivity could be switched off and on in a reversible manner by fully removing and then restoring PrP^Sc^ sialylation, respectively (59,60). In animals, the most pronounced activation of microglia was observed in brain regions with the lowest sialylation status of PrP^Sc^ (61). These findings strongly suggest a causal relationship between PrP^Sc^ sialylation status and microglia response. In the current study, in contrast to PrP^Sc^ dynamics, the intensity of microglia activation continues to increase after the third passage in a new host. This result supports the hypothesis that a certain level of PrP^Sc^-induced neuroinflammation is necessary for the disease to progress to a terminal stage. Since hyposialylated PrP^Sc^ is more potent than hypersialylated PrP^Sc^ in triggering neuroinflammation, this work suggests that the disease pathogenesis is driven not only by the quantity of PrP^Sc^ but also by its sialylation status.

Galactose residues at the terminal positions of N-linked glycans are anticipated to trigger ‘eat me’ phagocytic signaling within microglia (54,55,57,58). In our study, mouse-adapted S05 exhibited small but readily detectable granular aggregates of PrP^Sc^. These aggregates were primarily associated with microglia. Instances of PrP^Sc^ puncta were infrequent in astrocytes and nearly absent in neurons. It is worth noting that we cannot precisely determine the extent to which PrP^Sc^ is deposited in the form of diffuse particles on the surfaces of neurons and/or astrocytes. Due to challenges in distinguishing between cell surface PrP^C^ and diffuse PrP^Sc^, we implemented image analysis to filter out low-intensity diffuse signals.

The preferential association of S05 PrP^Sc^ with microglia stands in contrast to the neuron- or astrocyte-specific associations observed with PrP^Sc^ in other mouse-adapted strains, such as RML, 22L, or ME7 (6). Intriguingly, in comparison to RML, 22L, or ME7, the mouse-adapted S05 strain exhibited the shortest incubation time to the disease. Another mouse-adapted strain of synthetic origin, SSLOW, also predominantly colocalized with microglia, and its incubation time was even shorter than that of S05 (62). Moreover, SSLOW elicited a much stronger neuroinflammatory response compared to that observed in RML-, 22L-, or ME7-infected animals (62–64). In conjunction with previous studies, the current research suggests that there are relationships between the sialylation status of PrP^Sc^, the extent of neuroinflammation, and the incubation time to disease. While the precise mechanistic details of these relationships await further elucidation in future studies, recent years have unveiled intricate cross-talk between microglia and astrocytes (65). The reactive states of both astrocytes and microglia appear to be mutually dependent and regulated through multiple signaling pathways (65,66). In recent studies, reactive astrocytes associated with prion diseases were found to cause neurotoxicity and a breakdown of the blood-brain barrier (BBB) (67,68). Furthermore, it was observed that the extent of astrocyte activation inversely correlates with the incubation time to disease, suggesting that neuroinflammation propels disease progression (63). It would be interesting to explore whether microglia also contribute to non-cell-autonomous neuronal death and whether microglia-induced toxicity targets neurons directly or is mediated by cross-talk with astrocytes. In support of the idea on microglia-induced neurotoxicity, recent findings showed that reactive microglia isolated from prion-infected mice significantly increased their phagocytic activity toward both normal and disease-associated synaptosomes, revealing a lack of selectivity towards disease-associated substrates (69).

What are the molecular factors responsible for accelerating the replication rate of PrP^Sc^ during the initial adaptation stage? Recent cryo-microscopy studies have unveiled significant topological distinctions between hamster and mouse strains (70–72). It is likely that these topological preferences are encoded by species-specific PrP sequences. In PrP^Sc^ amyloid fibrils, each PrP molecule forms a single rung, taking on a V-shaped structure with two distinct N- and C-terminal lobes (70–72). In the hamster strain 263K, the groove between the N- and C-terminal lobes is notably wide (70), while in mouse strains (RML and ME7), the grooves are comparatively narrow (71,72). The N180 glycan (in mouse PrP) or N181 glycan (in hamster PrP) occupies the groove between the N- and C-terminal lobes, whereas N196 (in mouse PrP) or N197 (in hamster PrP) extends outward from the C-lobe (70–72). In hamster PrP^Sc^, the lobe topology allows for the accommodation of bulky hypersialylated N-glycans in each rung, i.e. by each PrP molecule (70). Since there are no constraints on N197 glycans either, hamster PrP^Sc^ is predominantly diglycosylated, reflecting the glycoform composition of hamster PrP^C^. Our hypothesis suggests that, in order to fit within the narrow groove and minimize electrostatic repulsion between negatively charged sialic acid residues of N180 glycans, diglycosylated PrP molecules must be partially excluded from mouse PrP^Sc^ fibrils. In mouse fibrils, diglycosylated PrP molecules have to alternate with monoglycosylated PrP molecules carrying a glycan at N197 or non-glycosylated molecules. In agreement with the above hypothesis, the removal of sialic acid residues from the PrP^C^ substrate was found to restore the glycoform ratio of the mouse strains to that of PrP^C^ (17).

Notably, in the first passage in mice, S05 exhibits the same glycoform ratio as hamster S05. This observation suggests that during this early passage, mouse PrP^C^ still propagates hamster-specific topology, albeit at the cost of a slower replication rate. Changes in the glycoform ratio throughout the strain adaptation process imply a shift towards a mouse-specific rung topology. It remains unclear whether this transformation occurs gradually, progressing through several distinct PrP^Sc^ states, or if it occurs in a single step, transitioning from hamster-specific to mouse-specific rung topology. If the latter is accurate, the alteration in glycoform ratio during several passages might be attributed to a gradual shift in the proportion of two PrP^Sc^ states, corresponding to hamster- and mouse-specific topologies. An alternative scenario suggests that the replication of PrP^Sc^ with hamster-specific rung topology in mice leads to the generation of multiple PrP^Sc^ variants through a deformed templating mechanism (48). While the majority of these variants may not propagate effectively in mouse PrP^C^, a variant featuring mouse-specific topology may eventually emerge through numerous trial-and-error seeding events (48). Regardless of the specific mechanism at play, it is anticipated that the replication rate of PrP^Sc^ will accelerate as a result of (i) the transition to mouse-specific rung topology and (ii) the switch to the selective recruitment of hyposialylated PrP^C^ molecules as a preferred substrate. Preferential recruitment of hyposialylated PrP^C^ molecules minimizes electrostatic repulsion between neighboring PrP molecules within PrP^Sc^ fibrils. In support of this mechanism, previous studies have demonstrated that the removal of sialic acid residues from PrP^C^ can significantly enhance PrP^Sc^ replication rates in sPMCAb, with increases ranging from 20- to 106-fold, depending on the strain (16,17).

Hamster-specific rung topology exhibits no preferences in recruiting sialoglycoforms, whereas mouse topologies display a pronounced selectivity in recruitment (17,18). Consequently, hamster PrP^Sc^ is predominantly diglycosylated, while PrP^Sc^ in mouse strains is predominantly monoglycosylated. If differences in the selectivity of PrP^C^ recruitment are indicative of PrP^Sc^ rung topology, one can extrapolate the rung topology to PrP^Sc^ associated with Creutzfeldt-Jakob diseases (CJDs), which exhibit distinct glycoform ratios (73,74). Predominantly diglycosylated PrP^Sc^ in variant CJD suggests a rung topology with a wider groove, whereas predominantly monoglycosylated PrP^Sc^ in sporadic CJD (sCJD) implies a topology with a narrower groove.

In summary, the current study illustrates that the adaptation of a prion strain to a new host is a multi-step process driven by several factors. During the initial phase, the replication of donor PrP^Sc^ using the PrP^C^ of a new host triggers alterations in the topology of PrP^Sc^. When the PrP^Sc^ topologies of the donor and recipient hosts are substantially different, the new topology influences the selectivity of recruitments on PrP^C^ sialoglycoforms. The selection of a different set of sialoglycoforms leads to changes in the decoration of PrP^Sc^ by carbohydrate moieties, ultimately resulting in alterations in the disease phenotype.

## Experimental procedures

### Preparation of 10% BH_s_

10% (wt/vol) of BH was prepared in PBS, pH 7.4, using glass/Teflon homogenizers attached to a cordless 12 V compact drill as previously described (75).

### Serial Protein Misfolding Cyclic Amplification with beads (sPMCAb)

As a source of S05 for sPMCAb, brain-derived materials from the 4^th^ serial passage of S05 in hamsters were used (49). 10% normal BH from healthy mice was prepared as described previously (76) and used as a substrate for sPMCAb (52). The standard sonication program consisted of 5 s sonication pulses at ∼200 watts applied every 10 min during a 24 h period, and the reactions were carried out in the presence of two 3/32” Teflon beads (McMaster-Carr, Robbinsville, NJ) (52). For each subsequent round, 20 μl of the reaction from the previous round was added to 80 μl of fresh substrate. To analyze the production of PrP^Sc^ in sPMCAb, 10 μl of each sample was supplemented with 5 μl of SDS and 5 μl of Proteinase K (PK) to the final concentrations of 0.25% and 50 μg/ml, respectively, followed by incubation at 37°C for 1 h. The digestion was terminated by the addition of SDS-sample buffer and heating the samples for 10 min in a boiling water bath.

### Serial transmission in mice

S05 and Mo S05 brain-derived materials were inoculated as 10% BH prepared as described above in PBS. Immediately before inoculation, each inoculum was further dispersed by 30 seconds of indirect sonication at ∼200 watts in a microplate horn of a sonicator (Qsonica, Newtown, CT). sPMCAb-derived material was diluted 10-fold in PBS supplemented with 1% (w/vol) bovine serum albumin before inoculation, to reduce the amount of detergent in the inoculum. Each C57BL/6 mouse received 20 μl of inoculum intracerebrally (IC) under 3% isoflurane anesthesia. After inoculation, animals were observed daily for signs of neurological disorders. Clinical signs included clasping hind legs, difficulty walking, abnormal gate, nesting problems and weight loss. The animals were euthanized when they were unable to rear and/or lost 20% of their weight.

### Antibodies

Primary antibodies used for immunoblotting and immunofluorescence were as follows: polyclonal anti-prion antibody ab3531 (#ab3531, Abcam, Cambridge, MA), prion protein monoclonal antibody clone SAF-84 (#189775, Cayman, Ann Arbor, MI), recombinant Anti-PrP Fab HuM-D18 (InPro Biotechnology, San Francisco, CA), polyclonal anti-Iba1 (#013-27691, FUJIFILM Wako Chemicals, Richmond, VA), polyclonal anti-GFAP (#AB5541, Millipore Sigma), polyclonal anti-MAP2 (#NB300-213, Novus Biologicals, Centennial, CO). The secondary antibodies used for immunoblotting were goat anti-rabbit IgG-HRP (#474-1506, KPL), goat anti-mouse IgG–HRP (#474-1806, KPL), and goat anti-human IgG, F(ab’)_2_ fragment specific, peroxidase-conjugated (#31414, Pierce, Rockford, IL). The secondary antibodies for immunofluorescence were goat anti-mouse IgG antibody (Alexa Fluor 546, #A-11003, ThermoFisher Scientific), donkey anti-rabbit IgG antibody (Alexa Fluor 488, #A-21206, ThermoFisher Scientific), goat anti-rabbit IgG antibody (Alexa Fluor 546, #A-11010, ThermoFisher Scientific), and goat anti-chicken IgG antibody (Alexa Fluor 488, #A-11039, ThermoFisher Scientific).

### PK digestion of BHs for Western blot

10% BHs were mixed with an equal volume of 4% sarcosyl in PBS, supplemented with 50 mM Tris, pH 7.5, digested with 20 μg/ml PK (New England BioLabs, Ipswich, MA) for 30 min at 37°C with 1000 rpm shaking using a DELFIA plate shaker (Perkin Elmer, Waltham, MA) and placed in a 37°C incubator. PK digestion was stopped by adding SDS sample buffer and heating the samples for 10 min in a boiling water bath. Samples were loaded onto NuPAGE 12% Bis-Tris gels, transferred to 0.22 μm PVDF membrane, and probed with anti-prion antibodies, as indicated.

### 2D electrophoresis

2D electrophoresis was performed as previously described (77). Briefly, 25 µL of samples digested with PK, supplemented with gel loading buffer and heated as described above, were solubilized for 1 h at room temperature in 200 µL solubilization buffer (8 M Urea, 2% (wt/vol) CHAPS, 5 mM TBP, 20 mM TrisHCl pH 8.0), then alkylated by adding 7 µL of 0.5 M iodoacetamide and incubated for 1 h at room temperature in the dark. Then, 1150 µL of ice-cold methanol was added and samples were incubated for 2 h at −20°C. After centrifugation at 16,000 g at 4°C, the supernatant was discarded, and the pellet was re-solubilized in 160 µL rehydration buffer (7 M urea, 2 M thiourea, 1% (wt/vol) DTT, 1% (wt/vol) CHAPS, 1% (wt/vol) Triton X-100, 1% (vol/vol) ampholyte, trace amount of Bromophenol Blue). Fixed immobilized pre-cast IPG strips (cat. # ZM0018, Life Technologies, Carlsbad, CA) with a linear pH gradient 3-10 were rehydrated in 155 µL of the resulting mixture overnight at room temperature inside IPG Runner cassettes (cat. # ZM0008, Life Technologies). Isoelectrofocusing (first dimension separation) was performed at room temperature with rising voltage (175 V for 15 minutes, then 175 – 2,000 V linear gradient for 45 minutes, then 2,000 V for 30 minutes) on Life Technologies Zoom Dual Power Supply using an XCell SureLock Mini-Cell Electrophoresis System (cat. # EI0001, Life Technologies). The IPG strips were then equilibrated for 15 minutes consecutively in (i) 6 M Urea, 20% (vol/vol) glycerol, 2% SDS, 375 mM Tris-HCl pH 8.8, 130 mM DTT and (ii) 6 M Urea, 20% (vol/vol) glycerol, 2% SDS, 375 mM Tris-HCl pH 8.8, 135 mM iodoacetamide, and loaded on 4 – 12% Bis-Tris ZOOM SDS-PAGE pre-cast gels (cat. # NP0330BOX, Life Technologies). For the second dimension, SDS-PAGE was performed for 1 h at 170 V. Immunoblotting was performed with D18 antibody.

### Western blot densitometry analysis

1D or 2D Western blot signals were visualized using FluorChem M imaging system (ProteinSimple, San Jose, CA). Densitometry was performed using AlphaView software (ProteinSimple). To generate individual sialylation profiles, 2D gels were rotated about 90°, to allow di-, mono- and non-glycosylated sets of spots to be defined as three vertical lanes using the “Lane Profile” function of AlphaView. Intensity profiles of di-, mono- and non-glycosylated “lanes” were imported to GraphPad for building the graphs shown in Figure 2A.

Individual sialoglycoforms were divided into two groups according to their positions on 2D Western blots. The position of the demarcation line for separating isoforms into hyper- and hyposialylated groups was determined based on previous work that employed a panel of sialidases for desialylating PrP^Sc^ and establishing a boundary between the two groups (77). Quantification of intensities of the five groups in 2D blots was done with the AlphaView “Multiplex band analysis” option. Within each 2D gel, two rectangular boxes of the same area were drawn around each group of spots (Figure 3C). The position of the rectangles was consistent between all 2D gels. The third rectangle of the same area was drawn on an empty part of the gel and used for background correction. The intensities of each set of spots were normalized by the sum of intensities in the whole region and plotted in GraphPad. 2D gels were run for three animals from each passage.

### Histopathology and Immunofluorescence

Formalin-fixed brains (sagittal or coronal 3 mm slices) were treated for 1 hour in 96% formic acid before being embedded in paraffin. 4 μm sections produced using a Leica RM2235 microtome (Leica Biosystems, Buffalo Grove, IL) were mounted on slides and processed for immunohistochemistry. Detection was performed by using AlexaFluor-488 and AlexaFluor-546 labeled secondary.

To expose epitopes, slides were subjected to 20 min of hydrated autoclaving at 121° C in trisodium citrate buffer, pH 6.0, with 0.05% Tween 20. For the detection of disease-associated PrP, an additional treatment in 88% formic acid was applied. PrP was stained with the anti-prion antibody SAF-84. To detect astrocytes, chicken anti-GFAP was used. For microglia and neurons, rabbit anti-Iba1 and chicken anti-MAP2 were used, respectively. An autofluorescence eliminator (#2160, Sigma-Aldrich, St. Louis, MO) and Image-iT Signal Enhancer (#11932S, Cell Signaling Technology, Danvers, MA) were used according to the original protocol to reduce background fluorescence. Fluorescent images were collected using an inverted microscope Nikon Eclipse TE2000-U (Nikon Instruments Inc, Melville, NY) equipped with an illumination system X-cite 120 (EXFO Photonics Solutions Inc., Exton, PA) and a cooled 12-bit CoolSnap HQ CCD camera (Photometrics, Tucson, AZ). Images were processed using WCIF ImageJ software (National Institute of Health, Bethesda, MD, United States).

### Statistics

Statistical analyses were performed using GraphPad Prism software, version 8.4.2 for Windows (GraphPad Software, San Diego, California USA). The box-and-whisker plot of incubation time to clinical disease was built using Excel. The midline denotes the median, the x represents the mean and the ends of the box-and-whisker plot denote the 25^th^ and 75^th^ percentiles. The whiskers extend from the ends of the box to the minimum value and maximum value.

### Study approval

The study was carried out in strict accordance with the recommendations in the Guide for the Care and Use of Laboratory Animals of the National Institutes of Health. The animal protocol was approved by the Institutional Animal Care and Use Committee of the University of Maryland, Baltimore (Assurance Number: A32000-01; Permit Number: 0215002).

## Data availability

All data are contained within the manuscript.

## Author contributions

**Olga Bocharova:** Investigation; Methodology; Formal analysis, Writing – Reviewing and Editing. **Natallia Makarava;** Conceptualization, Investigation; Formal analysis, Writing – Reviewing and Editing. **Narayan P. Pandit:** Investigation. **Kara Molesworth:** Resources, Investigation**. Ilia V. Baskakov:** Conceptualization, Funding acquisition, Project administration, Supervision, Writing – Original Draft.

## Funding and additional information

Financial support for this study was provided by National Institute of Health Grants R01 NS045585 and R01 NS129502 to IVB.

